# Generation of functional human adipose tissue in mice from primed progenitor cells

**DOI:** 10.1101/267427

**Authors:** Raziel Rojas-Rodriguez, Jorge Lujan-Hernandez, So Yun Min, Tiffany DeSouza, Patrick Teebagy, Anand Desai, Heather Tessier, Robert Slamin, Leah Siegel-Reamer, Cara Berg, Angel Baez, Janice Lalikos, Silvia Corvera

**Author notes:** Equal contributions (Raziel Rojas-Rodriguez, Jorge Lujan-Hernandez, So Yun Min, Tiffany DeSouza). Co-senior authors (Janice Lalikos, Silvia Corvera). Corresponding Author: Silvia Corvera, Program in Molecular Medicine, University of Massachusetts Medical School, 373 Plantation Street, Worcester MA 016581.

## Abstract

Adipose tissue is used extensively in reconstructive and regenerative therapies, but transplanted fat often undergoes inflammation and cell death, requiring further revision surgery. We report that functional human adipose tissue can be generated from mesenchymal progenitor cells in-vivo, providing an alternative approach to its therapeutic use. We leveraged previous findings that progenitor cells within the vasculature of human adipose tissue robustly proliferate in 3-dimensional culture under proangiogenic conditions. Implantation of these progenitor cells into immunocompromised mice results in differentiation towards non-adipocyte fates, incapable of generating a distinct tissue structure. However, priming of these progenitor cells in-vitro towards adipogenic differentiation results in formation of functional adipose tissue in-vivo. Mechanistically, priming induces the expression of genes encoding specific extracellular matrix and remodeling proteins, and induces extensive vascularization by host blood vessels. In comparison, grafts from adipose tissue obtained by liposuction undergo poor vascularization, adipocyte death, cyst formation, calcification and inefficient adiponectin secretion. Thus, primed mesenchymal adipose tissue progenitors reveal mechanisms of human adipose tissue development, and have potential to improve outcomes in reconstructive and regenerative medicine.

## Introduction

Adipose tissue (AT) is unique in its capacity to greatly expand as a function of nutrient consumption. Indeed, adipose tissue can expand to comprise up to 70% of human body mass (1). The capacity of AT to grow and promote angiogenesis has fueled the use of fat grafting in numerous medical contexts. (2-6). In some examples, AT is used for volume restoration after surgery for breast cancer (7), for soft tissue reconstruction in acquired or congenital malformations (8, 9) and to fill defects such as cranial fractures or fistulas in cranial neurosurgery (10-12). In the context of regeneration, mesenchymal cells from adipose tissue have been used for numerous purposes including cardiomyocyte (13), bowel (14), tendon and bone regeneration (15). In the setting of volume restoration, reabsorption and necrosis of grafted fat are a frequent complication (16, 17), resulting in the need for repeated surgical procedures in > 25% of cases. While empirical methods to improve fat grafting outcomes have been identified (18), a further understanding of the cellular and molecular mechanisms of adult human adipose tissue development are required to fully leverage its therapeutic potential.

Human adult AT is enriched in mesenchymal stem cells (ADSCs) capable of differentiation toward the adipogenic, myogenic, chondrogenic and osteogenic lineages (19-21). ADSCs may underlie the capacity of AT to expand, and play an important role in the outcome of AT grafting. Indeed, in some studies supplementation of engrafted AT with cells from the stromovascular fraction (SVF), which contains ADSCs, displayed significant improvements in volume retention (22), and controlled clinical trials support a specific role for ADSCs in graft improvement (23). The mechanisms by which ADSCs improve grafting are unclear, but they could function by providing new adipocytes or their vascular support (24, 25). ADSCs are obtained from the SVF of adipose tissue by a procedure that entails enzymatic or mechanical disruption of AT, followed by selection of adherent cells on plastic. Relatively few cells can be obtained by this procedure, and they display loss of differentiation potential with passaging (26, 27). Thus, the therapeutic use of ADSCs is restricted to instances in which relatively large amounts of adipose tissue can be collected.

A potential alternative approach to obtaining adipose tissue progenitor cells is to leverage the mechanisms by which adipose tissue expands physiologically. Previous studies indicate that development of adipose tissue is preceded by the formation of a vascular plexus, and lineage-tracing studies have shown that adipocyte progenitors are localized within the adult adipose tissue vasculature (28-36). These findings suggested the possibility that vascular expansion would be accompanied by adipocyte progenitor proliferation. We tested this hypothesis by culturing human adipose tissue explants under pro-angiogenic conditions that elicited capillary sprout formation, and found concomitant robust proliferation of adipocyte progenitor cells (37, 38). Moreover, adipocytes differentiated from these progenitors affected glucose metabolism when implanted into immunocompromised mice (38). We now wish to further understand whether these cells (referred to as Primed ADipocyte progenitor cells = PADS) are capable of forming structured, long-lived adipose tissue that could be used in surgical applications. In this manuscript, we use quantitative morphometric and biochemical methods to investigate the process by which PADS form structured, functional adipose tissue in-vivo, and compare it to conventional AT grafting. We also compare the molecular features of PADS to those of ADSCs prepared from the stromo-vascular fraction, and report the results of comparative gene expression analyses that identify unexpected pathways and genes associated with the efficient formation of functional AT in-vivo.

## Results

In vivo, human adipocytes contain large lipid droplets, display insulin responsive hypertrophy and leptin secretion, and display long-term survival. To determine the extent to which PADS represent functional human adipocytes we quantified mean adipocyte size over a long time in culture in the absence or presence of insulin. We find that PADS survive many weeks in culture, during which they transition from a multilocular to a unilocular phenotype (Figure 1A) and progressively increase in size. At 14 weeks after differentiation, cells are mostly unilocular, with abundant, functional mitochondria as seen by staining with the mitochondrial dye Mitotracker Red (Figure 1B). Insulin promotes adipocyte hypertrophy (Figure 1C), and stimulates leptin secretion (Figure 1D), indicating that PADS maintain the morphological and functional properties of human adipocytes.

**Figure 1.**
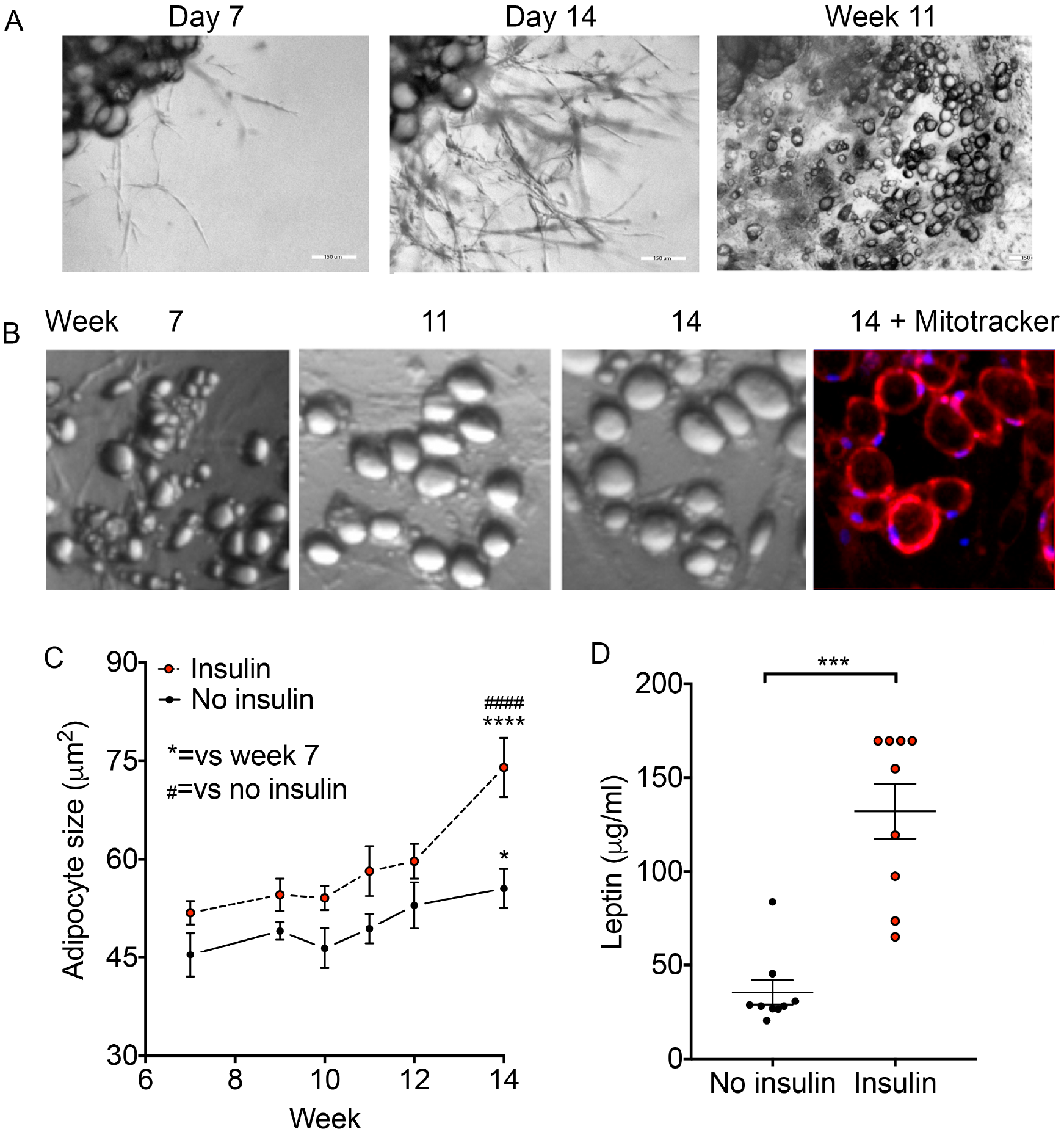
Development and properties of primed adipose progenitor cells (PADS) from human adipose tissue explants. **A.** Representative image of an explant from human subcutaneous adipose tissue (top left corner of each panel) in culturefor the time indicated above each panel. **B.** Higher magnification of specific section of well, taken at the times indicated. At week 14 Mitotracker red was added and cellsimaged after 30 min. **C.** Mean adipocyte size over time, fromexplants grown in the presence of absence of additional insulin added to theculture medium (100nM). Symbols are the means and bars SEM of adipocyte sizes, obtained from 5 independent wells per time point as described in Supplementary Figure 1. Statistical significance was estimated using 2-way ANOVA with Dunnet’s correction for multiple comparisons. *=p<0.05, ****= p < 0.0001 versus week 7; ####= p < 0.0001 versus no insulin. **D**. Concentration of leptin in media after 24h of culture at week 14. Shown are means and SEM of 8-10 wells. Statistical significance was estimated using two-tailed unpaired Mann-Whitney test. ***=p<0.001.

We then used morphometric and biochemical approaches to quantitatively assess the process by which PADS form adipose tissue in-vivo, and compared it with mature adipose tissue grafting. For the latter, a dry liposuction was performed on an excised specimen containing skin and subcutaneous tissue immediately following paniculectomy surgery. Liposuction fragments were then injected bilaterally under the skin of nude mice. In parallel, PADS suspended in Matrigel were injected unilaterally into a separate cohort of mice. Micro CT scans taken after 8 weeks (Figure 2A) revealed areas of low attenuation at the sites of injection of adipose tissue fragments (Figure 2A, bottom panel), and smaller, higher attenuated areas at sites of injection of PADS (Figure 2A, top panel). To measure graft functionality, we used a human-specific ELISA to measure circulating adiponectin, thereby reflecting viability and vascularization of grafted material. Human adiponectin was detectable in most engrafted mice (Figure 2B), but remarkably the amount of adiponectin per unit graft volume was 5-10 times higher in PADS compared to liposuction (Figure 2C). To better understand the differences in graft functionality we performed immunohistochemistry on the excised grafts. Visual inspection (Figure 2D) and quantitative analysis (Figure 2E) of the images revealed large de-cellularized regions in all grafts formed from liposuction tissue, probably corresponding to lipid-filled cysts, which are a frequent complication of fat grafting. These areas were undetectable in grafts formed from PADS (Figure 2D). These results indicate that PADS are capable of further differentiation and growth in vivo, and of efficient functional integration into the host.

**Figure 2.**
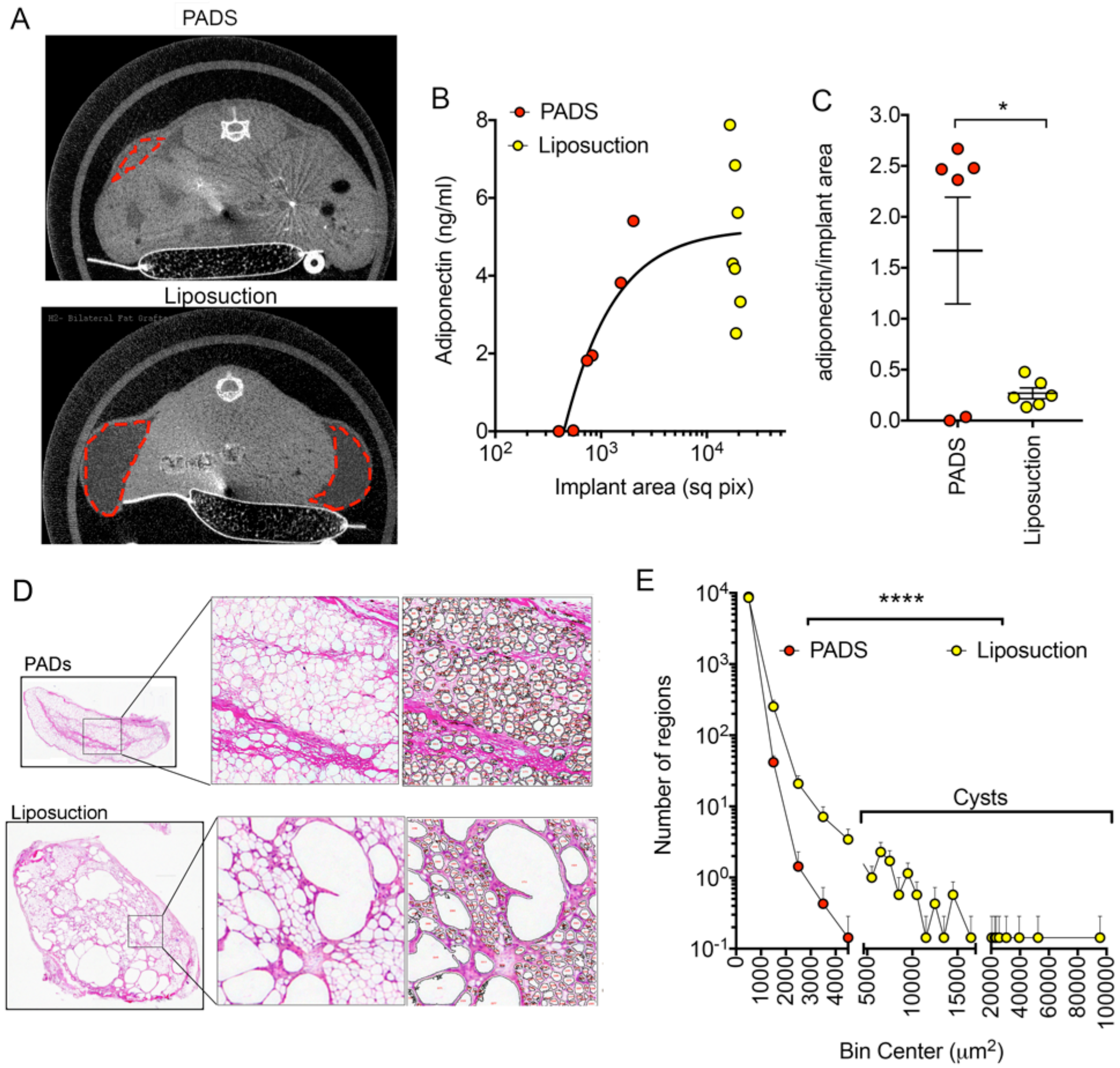
Comparison of grafts formed from PADS and from liposuction tissue. **A.** MicroCT scan of mice implanted with PADS (top panel) or liposuction tissue (bottom panel) after 23 or 7 weeks of implantation, respectively. Areas of low attenuation are enclosed by red dotted lines. **B.** Levels of human adiponectin in plasma from mice implanted as in A. Each symbol represents the valuefor each mouse in the cohort. **C.** Ratio of implant area versus circulating adiponectin. Each symbol represents one mouse, and lines represent the mean and SEM. Statistical significance was estimated using a two-tailed unpaired t-test with Welch’s correction for unequal standard deviations. * = p < 0.05. **D.** Histochemical staining of implants formed from PADS (top panels) orliposuction tissue (bottom panels). Expanded regions illustrate the algorithm output for selection of particles used for quantification. **E.** Histogram of particle sizes obtained from the mean values of 5 independent grafts from PADS and 7 independent grafts from liposuction tissue. Visual inspection was used to estimate adipocytes as those particles with values lower than between 100 and 5000 μ,m^2^. Statistical significance was estimated using the Friedman test of frequency distributions as implemented in Prism 7.0 ****=p<0.0001.

The mechanisms by which PADS integrate so efficiently into the host could involve recruitment of some of the human non-adipocyte progenitors to form functional vasculature (24). To test this hypothesis we measured the relative abundance of adipocyte and endothelial cells from human or mouse origin using species-specific qRT-PCR primers to the adipocyte lipid droplet marker perilipin 1 (mouse *Plin1,* human *PLIN1*), and to the mature endothelial cell marker VE-cadherin (mouse *Cdh5*, human *CDH5*). While grafts formed from PADS contained equal amount of human *PLIN1* compared to human subcutaneous adipose tissue (Figure 3A), grafts formed from liposuction contained less than half, reflecting loss of functional adipocytes (Figure 3A). This loss of adipocyte content occurred in spite of both types of grafts inducing vascularization from the host, as estimated by their high levels of mouse *Cdh5* (Figure 3B). Interestingly, grafts formed from liposuction tissue contained higher levels of human *CDH5* compared to control human adipose tissue, suggesting that human endothelial cells proliferate within the graft, possibly as a result of local hypoxia (Figure 3C). In contrast, no human *CDH5* was detectable in grafts formed from PADS, indicating that progenitor cells from the donor do not differentiate into endothelial cells under these conditions. Mouse *Plin1* was very low in both graft types, indicating negligible contamination from the host adipose tissue (Figure 3D). These results indicate that grafts formed from PADS are composed of human adipocytes perfused by mouse vasculature. To verify these results, we stained whole-mounts of the grafts with fluorescent lectins. Grafts formed from PADS were stained with mouse-specific lectin, but not with human lectin, confirming that the vascularization was derived from the mouse host and not the grafted cells (Figure 3E).

**Figure 3.**
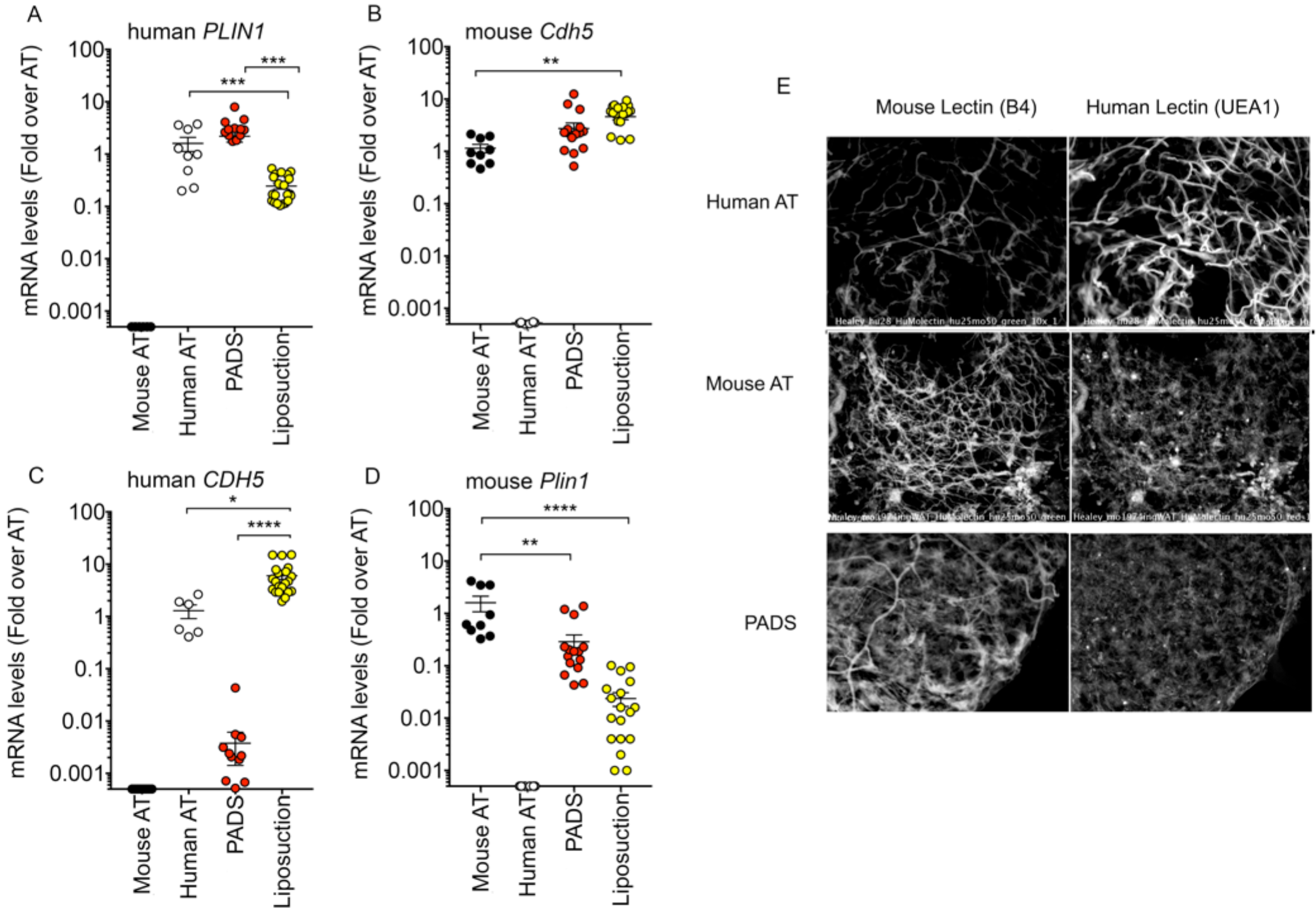
Human or mouse adipocyte and endothelial cell content in grafts formed from PADS and from liposuction tissue. A-D. RNA from excised grafts was analyzed using species-specific RT-PCR probes for the genes indicated above each panel. Symbols represents values for 3 tissue samples and 6-7 grafts per condition assayed in triplicate, and the mean and SEM of all grafts is shown. Statistical significance was estimated using multiple t-tests corrected for multiple comparisons using the Holm-Sidak method. *=p<0.05, **=p<0.01, ***=p<0.001, ****=p<0.00001. **E.** Whole mount staining of grafts formed from PADS usingmouse or human-specific lectins. Controls using human and mouse subcutaneousadipose tissue are illustrated.

The differences between the relative efficiency of engraftment of PADS compared to liposuction described above could be influenced by the large difference in volume between the two types of grafts, or by the use of Matrigel during grafting of PADS. The small volume of PADS compared to liposuction may mitigate hypoxia, thereby enabling adipocyte survival and development, and Matrigel has autonomous properties that facilitate adipogenesis (39). To better compare the inherent properties of PADS and liposuction tissue in the context of grafting, we performed experiments using smaller volumes of liposuction tissue with the inclusion of Matrigel, and also examined whether PADS would affect the engraftment of liposuction tissue. Three cohorts of mice were randomized to be injected unilaterally subcutaneously with 500μl of materials as follows: group A –liposuction tissue (250μl) in Matrigel (250μl); group B –PADS (100μl, ~10^7^ cells) in Matrigel (400μl); and group C –liposuction tissue (225 μl), PADS (50μl, ~5×10^6^ cells) in Matrigel (225μl). Five days following implantation micro CT scans revealed areas of low attenuation in all mice where liposuction fragments were engrafted (Figure 4, left top panels), but the regions where PADS were implanted were not distinguishable, probably due to their small size and content of lipid at the time of grafting (Figure 4 left middle panels). After 11 and 16 weeks, grafts formed from liposuction fragments were still clearly detectable, but had reduced in volume and developed small regions of high attenuation consistent with calcifications (Figure 4A, right panels, asterisks). In contrast, regions where PADS were implanted were now less attenuated, consistent with further differentiation and lipid accumulation of implanted cells, and no evidence of calcification was seen. At week 16 mice were sacrificed and grafts collected for further analysis. Grafts containing liposuction fragments were encapsulated, yellow, and surrounded by small blood vessels, whereas PADS were visible as non-encapsulated regions of adipose tissue under the skin resembling normal AT (Figure 4B). Histochemical findings were consistent with the micro CT findings, as grafts containing liposuction fragments displayed basophilic structures characteristic of calcification, and large de-cellularized regions, but grafts formed from PADS appeared as a well-vascularized adipose tissue parenchyma distinguishable from the mouse dermal adipose tissue (Figure 4C).

**Figure 4.**
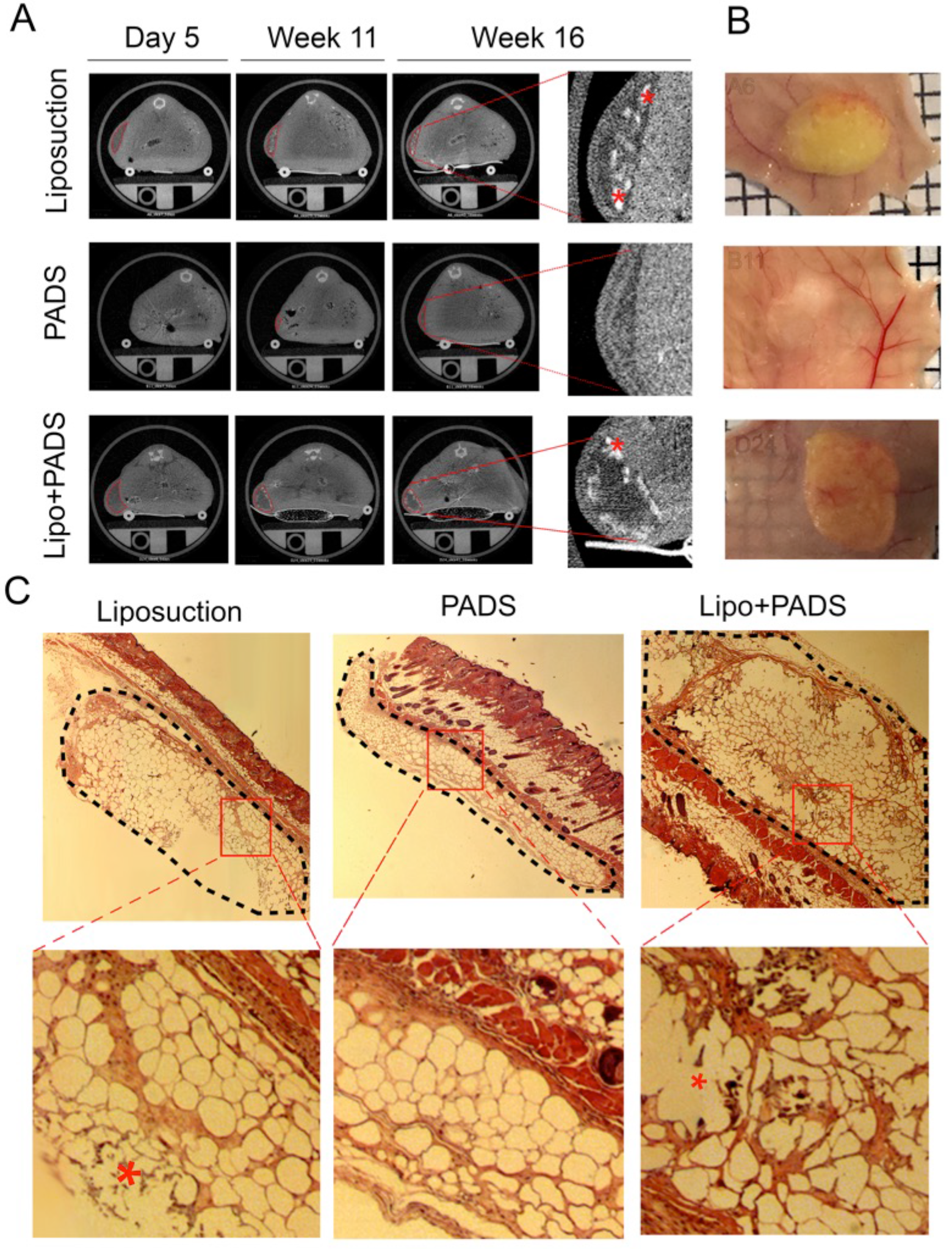
Development of grafts from PADS and liposuction tissue using comparable volumes. **A.** MicroCT scans of mice implanted with liposuction tissue (top panels) PADS (middle panels) and liposuction tissue supplemented with PADS (bottom panels) at the times following grafting indicated on top. Areas of low attenuationare outlined with red dottedtraces, and examples of high attenuation objectswithin these areas, possibly corresponding to calcifications, are indicated with asterisks. **B.** Macroscopic appearance of grafts at sacrifice. **C.** Histochemical staining of grafts. Grafted tissue under the skin is outlined with black, segmented traces, and basophilic objects within de-cellularized regions are illustrated with asterisks in the expanded images. Examples are representative of 7 grafts per condition.

Quantitative analysis of graft volume over time showed a decreasing trend in grafts formed by liposuction fragments. In contrast, the volume of grafts formed from PADS progressively increased (Figure 5A). Functionally, circulating adiponectin significantly increased between weeks 11 and 16 in mice harboring grafts from PADS, but not in mice harboring liposuction tissue (Figure 5B). Moreover, grafts formed from PADS were much more efficiently integrated, producing over 10x more adiponectin relative to graft volume at 16 weeks after implantation (Figure 5C). These results are consistent with those seen using larger graft volumes (Figure 2), but are more pronounced when volumes of PADS and liposuction tissue are approximated. This is clearly appreciated through analysis of both cohorts using a three-component agonist vs. response fit (Figure 5D), which illustrates that the slope of graft volume to adiponectin production is much steeper for grafts formed from PADS. Therefore, the enhanced ability of PADS to produce adiponectin cannot be attributed to a smaller graft volume, but to an inherent difference in the ability of adipocytes to survive and differentiate.

**Figure 5.**
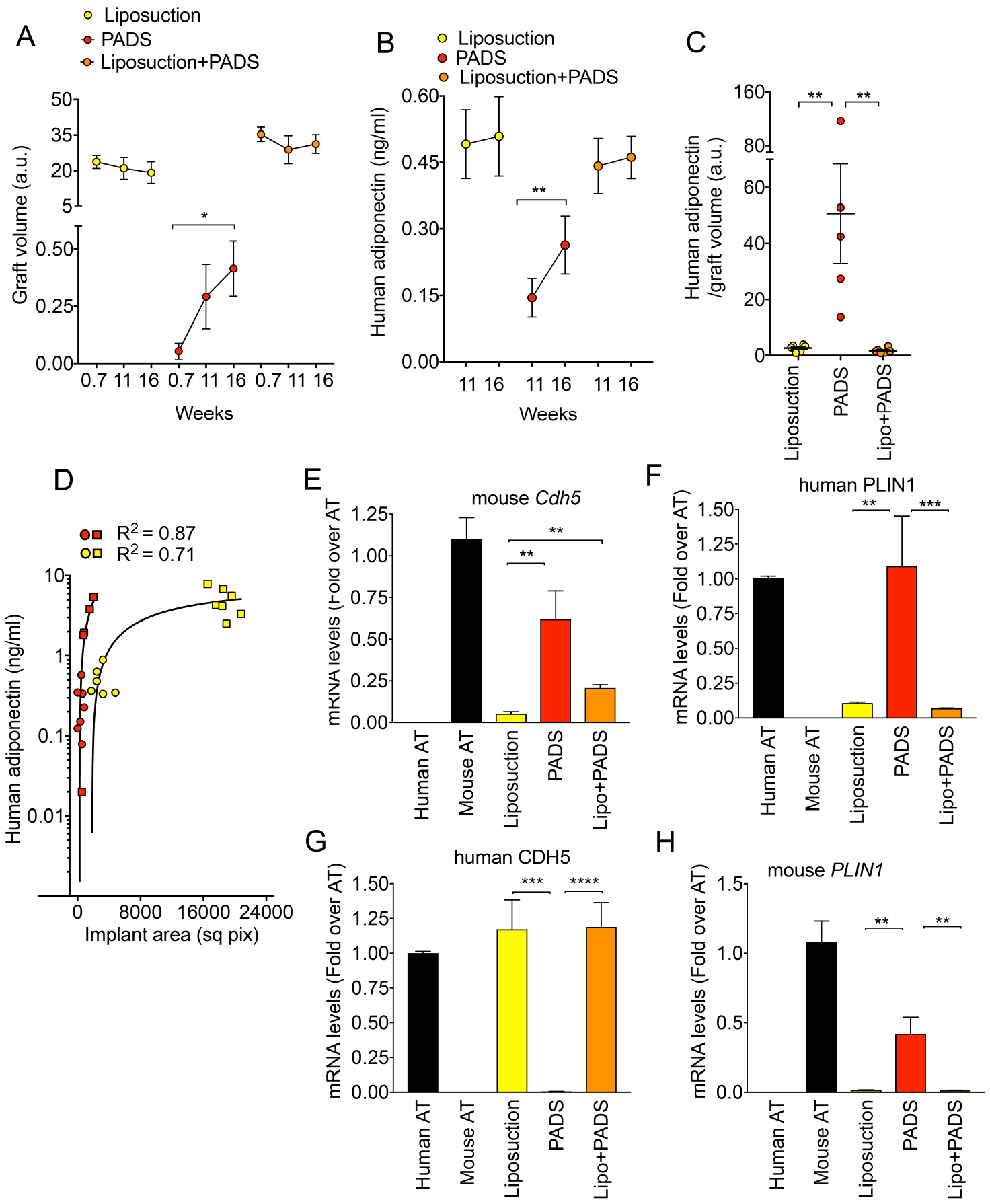
Quantification of graft volume, functionality and endothelial or adipose cell content. **A.** Means and SEM of graft volumes calculatedfrom microCT scans, and **B.** Means and SEM of human adiponectin in plasma,obtained at the times shown in the x-axes, from 7 mice per conditions implanted with liposuction tissue, PADS, or liposuction tissue supplemented by PADS. Statistical significance was estimated using paired t-tests. *=p<0.05, **=p<0.01. **C.** Means and SEM of the ratios of human adiponectin to graft volumes at 16 weeks post grafting in mice harboring grafts from liposuction tissue, PADS, or liposuction tissue supplemented by PADS. Statistical significance was estimated using one way ANOVA with Tukey’s correction for multiple comparisons, **=p<0.01. **D.** Relationship between graft volume and plasma adiponectin in mice from cohorts 1 (squares) and 2 (circles). Each symbol corresponds to values from one mouse. **E-H.** RNA from excised grafts was analyzed using species-specific RT-PCR probes for the genes indicated above each panel. Bars represent the mean, and lines the SEM of all grafts. Statistical significance was estimated using the Krustal-Wallis test with Dunn’s correction for multiple comparisons **=p<0.01, ***=p<0.001, ****=p<0.00001.

Vascularization by mouse blood vessels as assessed by mouse *Cdh5* mRNA expression, was lower in this cohort compared to that seen with larger volumes, suggesting that a smaller graft volume elicits overall lower angiogenic stimulation, possibly as a consequence of lessened hypoxia. Interestingly, grafts formed from liposuction tissue supplemented by PADS induced more vascularization compared to liposuction tissue alone, demonstrating a significant pro-angiogenic effect of these cells (Figure 5E). Nevertheless, grafts from liposuction tissue still displayed a large loss of adipocyte viability as estimated by low expression of human *PLIN1* (Figure 5F), despite maintenance of human *CDH5* (Figure 5G). Interestingly, a substantial amount of mouse *Plin1* was detected in PADS (Figure 5H); this may represent mouse AT contamination, or newly forming adipocytes from progenitors carried into the graft by mouse vasculature. This former seems unlikely, as mouse adipose tissue would be expected contaminate all grafts equally. These results indicate that PADS can form functional adipose tissue, by promoting continued growth and differentiation of adipocytes and eliciting vascularization.

To better understand the mechanisms whereby PADS can form functional adipose tissue, we examined the ability of different types of progenitors to form adipose tissue in-vivo. We compared PADS before differentiation (pre-PADS), ADSCs derived from the stromovascular fraction obtained by standardized procedures, and differentiated PADS (Figure 6A). We first conducted RT-PCR to determine levels of a broad panel of surface markers (Figure 6B). Pre-PADS displayed significantly higher levels of numerous markers, including PDGFRα, which has been associated with adipocyte progenitors in mouse models. In mice implanted with either ADSCs or pre-PADS no circulating human adiponectin could be detected up to 17 weeks post-implantation, indicating that these cells cannot form a functional adipose tissue graft. Fluorescent lectin staining of excised Matrigel remnants revealed the presence of numerous cell nuclei, but no lipid droplet accumulation where ADSCs were implanted (Figure 6C). In contrast, remnants from pre-PADS displayed some lipid droplets. Interestingly, human-specific lectin stained vessel-like structures in remnants from ADSCs (Figure 6C, arrows), but not in remnants from pre-PADS, suggesting that ADSCs are capable of forming micro vessels in vivo. Results from RT-PCR of the remnants (Figure 6D) were consistent with this possibility, as ADSC remnants expressed significantly higher levels of human *CDH5* compared with either pre-PADS or PADS. The levels of expression of human *PLIN1* were also consistent with morphological evidence for adipose differentiation of PADS, followed by pre-PADS but not of ADSCs in-vivo. The levels of mouse *Cdh5* indicate that pre-PADS and ADSCs induced host vascularization poorly compared to PADS. These experiments indicate that PADS and ADSCs differ in that, in-vivo, PADS can form adipocytes but no vascular structures from donor-derived cells, while ADSCs are able to form donor-derived vascular structures.

**Figure 6.**
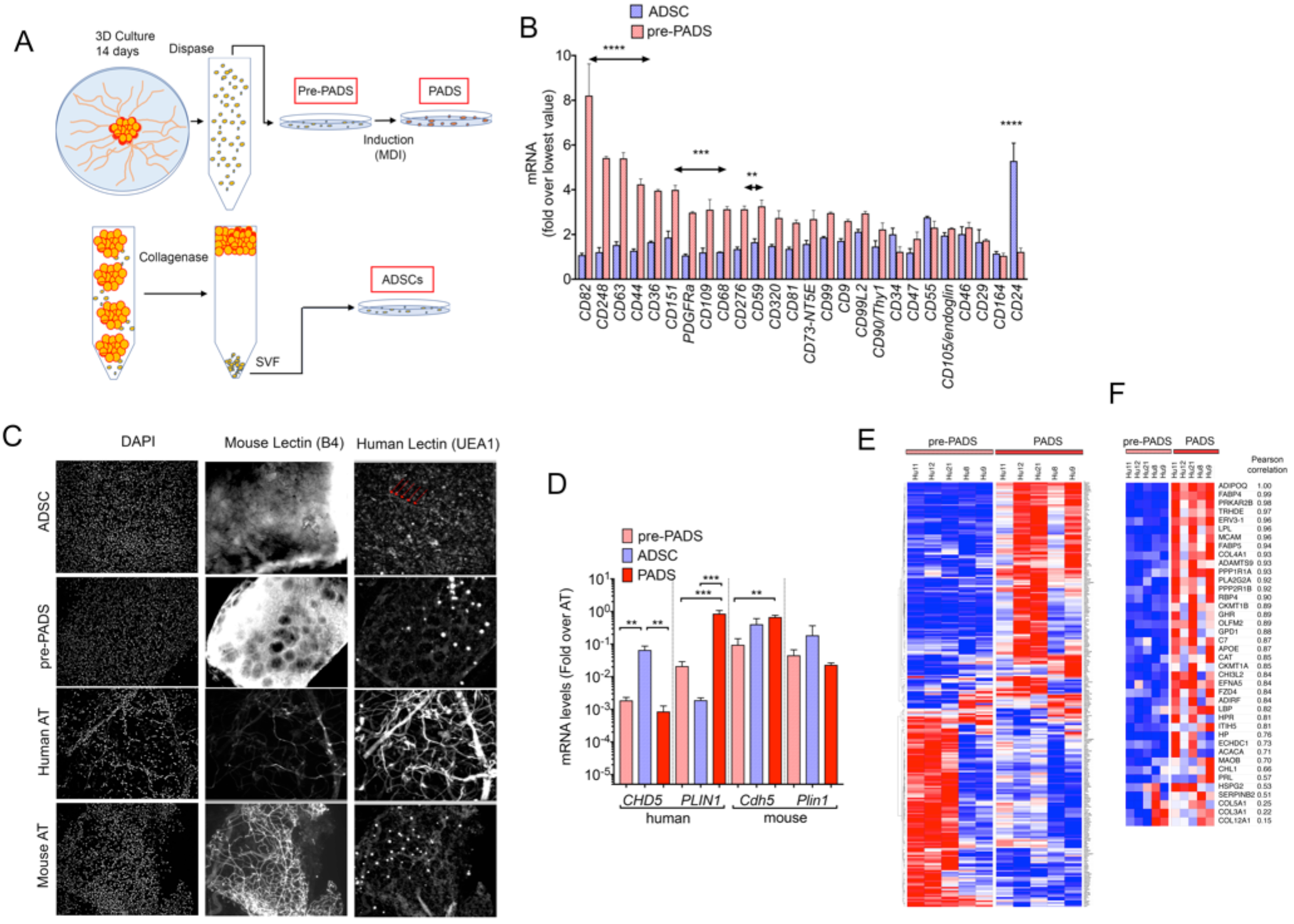
**A.** Schematic representing the methods to obtain pre-PADS, PADS and ADSCs. **B.** RNA from two independent preparations of ADSCs and pre-PADS was probed for the mesenchymal stem cell markers depicted in the x-axis. Values are expressed as the fold over the lowest value for each probe set. Statistical significance was estimated using multiple t-tests corrected for multiple comparisons usingthe Holm-Sidak method. Double headed arrows are placed over sets of probes that differed significantly, at the levels indicated **=p<0.01, ***=p<0.001, ****=p<0.00001. **C.** Whole mount staining of grafts formed from ADSCs and pre-PADS using DAPI, and mouse or human specific lectins. Control human and mouse adipose tissue are included for comparison. Linear structures stained with human-specific lectin in grafts formed from ADSCs are indicated with red arrows. **D.** RNA from excised grafts was analyzed using species-specific RT-PCR probes for the genes indicated in the x-axis. Bars represent the mean, and lines the SEM of 7 grafts per condition assayed in duplicate. Statistical significance was estimated using the Krustal-Wallis test with Dunn’s correction for multiple comparisons **=p<0.01, ***=p<0.001, ****=p<0.00001. **E.** Heat map of differentially expressed genes between pre-PADS and PADS, obtained from AT from 5 different individuals. **F.** Heat map of differentially expressed genes containing the GO term“extracellular”.

Because differentiated PADS were significantly more capable of forming functional adipose tissue in-vivo compared to pre-PADS, we performed a gene array comparison to identify those genes that promote in-vivo adipogenesis. Of >33,000 mRNAs detected by the HTA-2 arrays, 269 coding genes were differentially expressed with an adjusted p-value of less than 0.05 (Figure 6E and Supplementary Table 1). Of these, 40 genes containing the GO cellular component term “extracellular” (including “exosomal”) were up-regulated in response to differentiation (Figure 6F and Table 1). The top biological processes identified through enrichment analysis of this set were “response to oxygen” and “response to wounding”, (Table 2), and included 4 collagen species (COL4A1, COL5A1, COL3A1 and COL12A1), adhesion proteins (MCAM/CD146,CHL1,HSPG2), extracellular matrix remodeling proteins (ADAMTS9,OLFM2,SERPINB2), and secretory hydrolases (PLA2G2A,CH3L2). Also in this list were the expected adipokines and adipocyte secreted proteins ADIPOQ, FABP4, LPL, FABP5 and RBP4. Interestingly, none of these genes corresponded to known classical pro-angiogenic growth factors suggesting that the mechanisms by which PADS are capable of forming functional adipose tissue in-vivo is through the generation of an extracellular matrix niche optimal for adipogenic differentiation and host vascularization.

**TABLE 1.**
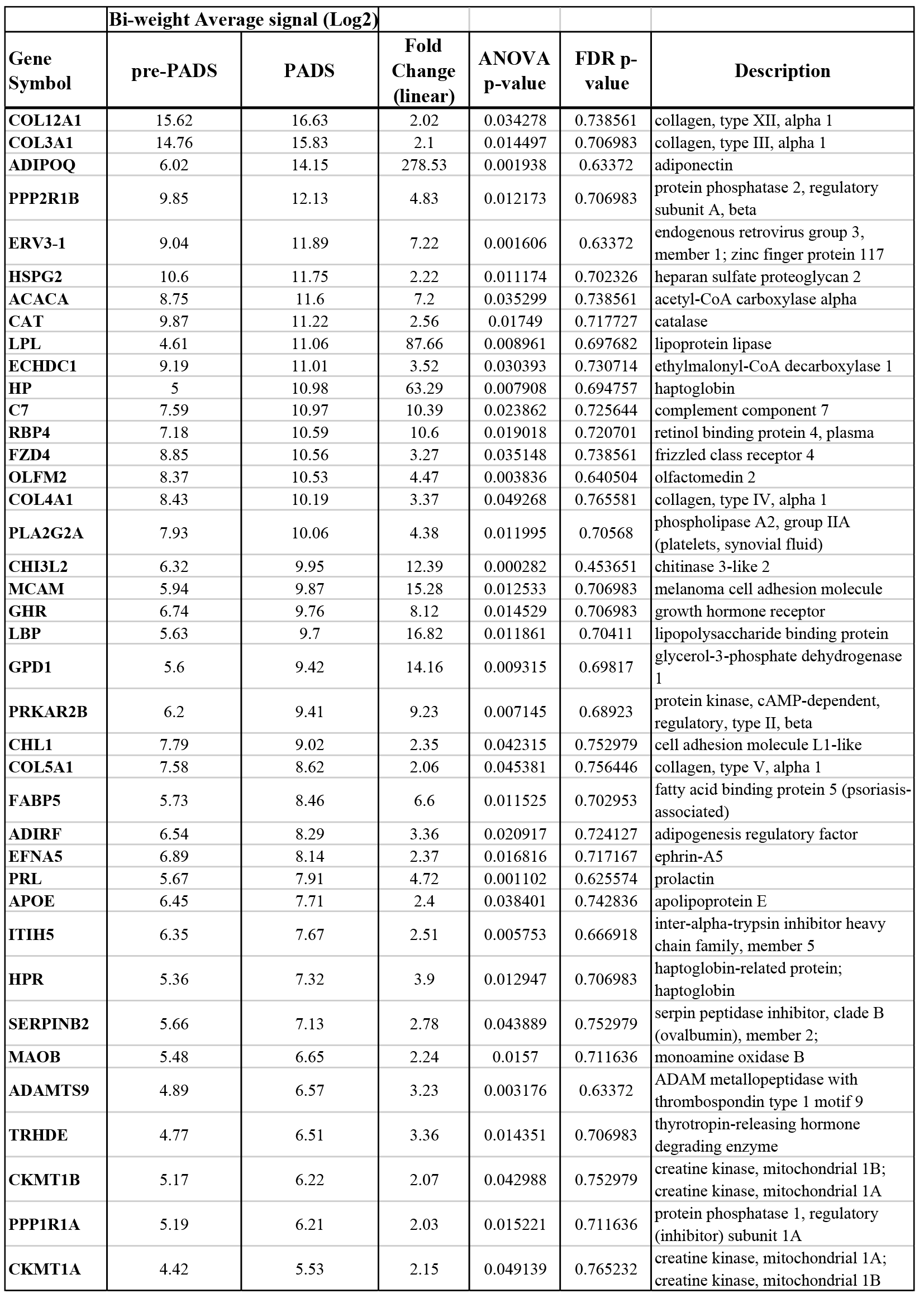
Genes containing the GO:cell component term “extracellular”that are upregulated in PADS compared to pre-PADS. The signals are means from 5 independent arrays, and values are sorted based on intensity of signal in PADS.

**TABLE 2.** Biological processes showing enrichment of genes containing the GO:cell component term “extracellular” that are upregulated in PADS compared to pre-PADS.

| GO Biological Process ID | Name | p-value | q-value Bonferroni | Hit Count in Query List | Hit Count in Genome | Hit in Query List |
| --- | --- | --- | --- | --- | --- | --- |
| GO:1901700 | response to oxygen-containing compound | 6.49E-09 | 1.23E-05 | 17 | 1614 | COL3A1,FZD4,COL4A1,GHR,GPD1,PRKAR2B,CAT,LBP,ADIPOQ,EFNA5,APOE,ACACA,MAOB,HP,PRL,LPL,RBP4 |
| GO:0009611 | response to wounding | 4.24E-08 | 8.06E-05 | 13 | 967 | CHL1,COL3A1,MCAM,PLA2G2A,PRKAR2B,COL5A1,LBP,ADIPOQ,C7,APOE,LPL,FABP5,SERPINB2 |
| GO:0044236 | multicellular organism metabolic process | 5.09E-07 | 9.68E-04 | 6 | 144 | COL3A1,COL4A1,GHR,COL5A1,COL12A1,ACACA |
| GO:0044255 | cellular lipid metabolic process | 1.06E-06 | 2.01E-03 | 12 | 1064 | GHR,GPD1,PLA2G2A,PRKAR2B,HSPG2,CAT,ADIPOQ,APOE,ACACA,LPL,FABP5,RBP4 |
| GO:0006600 | creatine metabolic | 1.39E-06 | 2.63E-03 | 3 | 11 | GHR,CKMT1B,CKMT1A |
| GO:0006641 | triglyceride metabolic process | 4.01E-06 | 7.62E-03 | 5 | 115 | GPD1,CAT,APOE,LPL,FABP5 |
| GO:1901701 | cellular response to oxygen-containing compound | 4.27E-06 | 8.12E-03 | 11 | 1001 | COL3A1,FZD4,COL4A1,GHR,GPD1,PRKAR2B,LBP,ADIPOQ,EFNA5,ACACA,PRL |
| GO:0044259 | multicellular organismal macromolecule | 6.52E-06 | 1.24E-02 | 5 | 127 | COL3A1,COL4A1,COL5A1,COL12A1,ACACA |
| GO:0006639 | acylglycerol metabolic process | 6.77E-06 | 1.29E-02 | 5 | 128 | GPD1,CAT,APOE,LPL,FABP5 |

## Discussion

The results shown in this manuscript contribute to our understanding of the mechanisms by which human adipose tissue develops in-vivo, and have practical implications for reconstructive and regenerative therapeutics. To our knowledge, this is the first method by which functional AT can be produced through in-vitro generated progenitors. The capacity of these progenitors to generate functional tissue in-vivo is related to their robust propagation in-vitro and to the priming of adipose differentiation prior to engraftment. These procedures enrich the progenitor population for cells that differentiate into mature adipocytes, and express genes for extracellular matrix components capable of supporting continued adipogenic differentiation and vascularization by host cells.

The limitations of this work are that PADS are heterogeneous and we cannot fully describe their complexity nor identify the specific complement of cells required for adipose tissue formation in-vivo. This next step will require the development of methods to identify and enrich the different progenitor subpopulations, and to examine their ability to engraft singly or in combinations. Nevertheless, the capacity of PADS to form functional adipose tissue has been reproduced in our laboratory with a minimum of 5 different cell preparations from different donors, using both NSG (38) and nude mouse strains (current results), indicating that robust conserved mechanisms are at play.

A remarkable feature of PADS is the number of cells that can be obtained by these methods. Standard methods to obtain ADSCs involve obtaining the stromovascular fraction of adipose tissue by collagenase digestion, and selection of mesenchymal progenitors through their capacity to attach to plastic surfaces. In contrast, PADS are obtained by placing non-digested tissue in 3-dimensional culture in the presence of pro-angiogenic basal media and growth factors. The reported number of cells and ADSCs that can be obtained from liposuction tissue varies widely, from 5×10^5^ to 2×10^6^ cells per gram (g) of adipose tissue, with 1 to 10% of these considered to be ADSCs (15, 40). While cells can be expanded in culture, they cease to replicate after 15 population doublings (41) and display in increasing senescence markers after 4 doublings (27). Based on these reports, the maximal yield of ADSCs from 1g of tissue can be estimated to be a maximum of 6 × 10^6^ after 5 passages. In contrast, explants from 0.25 cm^3^ (~200 mg) of adipose tissue yielded 2 × 10^7^ cells after 14 days in culture, and 8 × 10^7^ adherent cells 72 h after plating in plastic. In our hands PADS are passaged in a 1:2 split after this step to yield 1.7 × 10^8^ cells, which are frozen and can be expanded through 5 population doublings with no decrease in adipose differentiation capacity. Therefore ~5×10^9^ PADS can be obtained from 1 g of human adipose tissue by the procedures described here, representing a 1000x higher yield than ADSCs. It is possible that 3-dimensional organ culture triggers a physiological induction of progenitor cell proliferation, which in combination with minimal damage from mechanical or chemical stress, leads to very high yields of progenitor cells.

The differences between ADSCs and PADS extend beyond the yield of cells obtained through different techniques, as cell surface marker content differs significantly (Figure 6B) as do functional outcomes. It is interesting that ADSCs seem more capable of producing vascular structures in-vivo (Figure 6C), consistent with numerous reports of their pro-angiogenic effects (42-47). In contrast, PADS are more able to differentiate into adipocytes, and can induce host angiogenesis. These functional differences are likely due to the different complement of cells making up the ADSC and PADS populations, or acquired differences in functionality stemming from the stark difference in the way they are obtained.

To better understand the mechanisms that are directly involved in the formation of adipose tissue in-vitro, we leveraged the finding that pre-PADS are much less able to form tissue compared to PADS. This comparison is most informative, as the only difference between these cells is the exposure to a minimal adipogenic cocktail of methylisobutylxanthine, insulin and dexamethasone for three days, followed by 7 days of culture in DMEM. This leads to a dramatic difference in-vivo, where PADS continue to differentiate, accumulate lipids, become vascularized and secrete adiponectin into the circulation, whereas pre-PADS fail to form functional tissue. Comparison of gene expression profiles between these cells reveals that the most significant biological pathways enriched in PADS are related to canonical adipose tissue function, with the top 20 corresponding to lipid metabolic processes including storage and catabolism. However, the first non-lipid related pathway significantly enriched is that of wound healing (Supplementary Table 2). When the differentially expressed genes are restricted to those associated with the extracellular compartment, the second most enriched pathway corresponds to wound healing (Table 2). Strikingly, none of the genes previously associated with angiogenesis in the context of adipose tissue mesenchymal stem cells, which include VEGF FGF-2, HGF and TGF-ß (48), were found. Rather, extracellular matrix components, including 4 different collagen types were the predominant associated genes. One of these, COL4A1 is essential for small vessel development and pathogenic mutations cause vascular abnormalities (49-52), suggesting it may have an important role in vascularization of PADS for engraftment. Also, mutations in COL3A1 give rise to vascular Ehlers-Danlos syndrome (53), indicating a role for this collagen type in vascular integrity. Other identified genes include MCAM/CD146, which has been implicated in lymphangiogenesis (54), brain endothelial cell functions (55) and angiogenesis (56, 57). The interactions among these genes may also be critically important for tissue development, as has been reported for the morphogenesis of vascular structures through interactions between ADAMTS9 with HSPG2 (58, 59). How these genes exert these specific actions is not known, but the use of PADS grafting as a model may be useful for elucidating their molecular functions.

The therapeutic use of PADS is an exciting prospect in reconstructive surgery, particularly in instances such as wound healing or restoration of soft tissue volume after cancer surgery. PADS could be used alone for small reconstructive requirements or in combination with liposuction tissue to enhance large volume grafting. In our study, supplementation of liposuction tissue with PADS significantly enhanced vascularization, albeit without effect on graft function as assessed by adiponectin secretion. Further experiments will determine whether an optimal ratio of PADS to tissue achieve functionally improved grafting, and will focus on the development of culture systems compliant with good manufacturing practices.

## Materials and Methods

### Specific reagents

Matrigel (BD biosciences); EGM-2 MV (Lonza); Adiponectin human-specific ELISA (Invitrogen KHP0041); Rhodamine Ulex europaeus Agglutinin 1 (Vector labs RL-1062); Isolectin GS IB4 Alexa Fluor 488 Conjugate (Life Technologies I21411).

### Adipose Tissue

Subcutaneous adipose tissue for grafting and for derivation of PADS was obtained from panniculectomies with no a-priori selection of individual donors. All specimens were collected in accordance with procedures approved by the University of Massachusetts Institutional Review Board.

### Liposuction tissue

Tissue was obtained from panniculectomies performed at our institution, under an approved IRB. Immediately after excision of the panniculus from the patient, adipose tissue was extracted manually in multiple passes with a 3mm cannula, and centrifuged at 3000 RPM for 3 minutes. Aqueous and lipid components were discarded and fat used for grafting directly, or after mixing with an equal volume of Matrigel by gentle inversion, as indicated in each experiment. In some experiments, the 50/50 mixture of liposuction tissue and Matrigel was supplemented with PADS as described in the results section.

### Cells

Detailed methods for harvesting adipose tissue, culture of adipose tissue explants in Matrigel, and harvesting of single cells from explant growth are published (60). In brief, explants from human subcutaneous adipose tissue were cultured in EBM-2 media supplemented with endothelial growth factors (EGM-2 MV) (Lonza) fr 14 days. Single cells suspensions from capillary growth were obtained using dispase (yield ~2 × 10^7^ cells from one 10 cm plate) and plated into two 15 cm tissue culture dish. After 72 h, cells were split 1:2, grown for an additional 72h, recovered by trypsinization (yield ~8×10^7^) and frozen into 10 vials. To generate PADS for grafting, single vials were thawed out and plated into 1 × 150mm dish per graft and grown to confluence in EGM-2 MV. Differentiation was induced by replacement of EGM-2 MV by DMEM + 10%FBS, 0.5 mM 3-isobutyl-1-methylxanthine, 1μM dexamethasone, and 1μg/ml insulin (MDI). 72h later, the differentiation medium was replaced by DMEM-FBS, which was replaced every 48 hours for 10 days.

### Mice

All animal use was in accordance with the guidelines of the Animal Care and Use Committee of the University of Massachusetts Medical School. Male nude mice were obtained from the Charles River Laboratories. Liposuction tissue, PADS or a combination as indicated were grafted subcutaneously into the flank 5 cm cephalad to the tail and 3 cm lateral to the spine. Under sterile conditions, a 18G IV cannula was used to penetrate the skin at the base of the tail of each mouse. The needle was withdrawn and the shield of the catheter was advanced to the selected area, creating a tunnel by blunt dissection. A semi-rigid circular-shaped plastic splint with central hole of 1.5 cm diameter was positioned on the skin encircling the selected area to be grafted to ensure proper location and prevent displacement of the graft at the time of injection. The splint was left in place for 3 days and then removed. Serial volumetric analysis by Micro-CT Scanning was performed as described (61). Immediately after euthanasia, grafts were dissected, and half of the graft was fixed in 4% paraformaldehyde for microscopy and half used for RNA extraction using TRIzol. For RT-PCR, total RNA was reverse-transcribed using iScript cDNA synthesis kit (Bio-Rad). cDNA was used as template for qRT-PCR using the iQ SYBR Green Supermix kit (Bio-Rad) and the CFX96 Real-Time System (Bio-Rad). Human ribosomal protein L4 (RPL4) was used for normalization.

### Gene expression

For RT-PCR total RNA was reverse-transcribed using the iScript cDNA synthesis kit (Bio-Rad). cDNA was used as template for qRT-PCR using the iQ SYBR Green Supermix kit (Bio-Rad) and the CFX96 Real-Time System (Bio-Rad). Ribosomal protein L4 (RPL4) was used for normalization. Human and mouse-specific probe sets are shown in Supplementary table 3. For Affymetrix arrays, total RNA was isolated using TRIzol, from pre-PADS and PADS obtained from 5 independent subjects. Affymetrix protocols were followed for the preparation of cRNA, which was hybridized to HTA-2.0 arrays. Raw expression data collected from an Affymetrix HP GeneArrayScanner was normalized across all data sets using the RMA algorithm as implemented by the Affymetrix Expression Console. Differential expression analysis was performed using the Affymetrix Transcriptome Analysis Console v.3.0. Pathway analysis of transcriptomic data was performed using ToppGene (62).

### Histochemistry and quantification

Samples were fixed in 4% formaldehyde and embedded in paraffin. Tissue sections (8-μm), were mounted on Superfrost Plus microscope slides (Fisher Scientific), and stained with hematoxylin and eosin (H&E). 10X images were taken using brightfield microscopy (Zeiss Axiovert 35) and AxioCam Icc 1 digital camera (Zeiss). Adipocyte (area) size was determined using an automated procedure based on the open software platform FIJI (63). In brief, images were imported, local contrast enhanced (CLAHE, blocksize=263, histogram=256, maximum=40), converted to 8-bit grayscale image, binarized (threshold 136, 255, dark background), eroded (iterations=1, count=6, black, pad do=Erode), and object number and size were measured (Analyze Particles, size=100-Infinity circularity=0.10-1.00). Frequency distributions were obtained for each section using a bin size of 1000, and the means and SEM for each bin were calculated to generate the composite histogram for each graft type. For whole mount staining tissue fragments were fixed in 4% formaldehyde, washed and stained with a mixture of 50 ug/ml each Rhodamine Ulex europaeus Agglutinin 1 (Vector labs RL-1062), and Isolectin GS IB4 Alexa Fluor 488 Conjugate (Life Technologies I21411) for 1hr at room temperature. After washing in PBS, fragments were mounted between 1.5mm coverslips sealed with Pro-Long Gold Antifade Reagent (Life Technologies).

### Statistical analysis

GraphPad Prism 7.0 was used for all analyses. Parametric or non-parametric test were chosen based on results from the D’Agostino-Pearson omnibus normality test, and are described in each figure.

## Acknowledgements

This work was supported by grant DK089101-04 to SC, and a Pilot Project grant from UL1 TR000161-05 to SC and JL.

